# Early life-stage thermal resilience is determined by climate-linked regulatory variation

**DOI:** 10.1101/2025.07.07.663603

**Authors:** Joaquin C. B. Nunez, Sumaetee Tangwancharoen, Kylie M. Finnegan, Eliza M. Bufferd, Olin C. King, Luke A. Proud, Brent L. Lockwood

**Author notes:** Equal contribution (Eliza M. Bufferd, Olin C. King). Dept. of Organismic and Evolutionary Biology, Harvard University, Cambridge, MA, USA.

## Abstract

Despite decades of research in environmental change, we know relatively little about the genetics of environmentally influenced traits across the life cycle of species with complex life histories. Previously, we reported that natural variation in heat tolerance is life-stage specific in *Drosophila melanogaster*, suggesting that thermal selection predominantly targets the early embryonic life stage. Here, we used advanced introgression and pooled whole-genome resequencing to map the genomic basis of enhanced embryonic heat tolerance in a neotropical line of *D. melanogaster*. We identified two loci on chromosomes 2R and X that were consistently targeted by 16 generations of thermal selection across six replicate introgressions. We compared alleles in these regions to published datasets of natural variation from North America and Europe using the DEST dataset. This analysis revealed that two SNPs associated with embryonic heat tolerance exhibited both clinal and seasonal patterns, with the seasonal variation significantly correlated with environmental variability in average precipitation and temperature variance across space and time. Further, tropical alleles at both loci exhibited enhanced embryonic heat tolerance in the *Drosophila* Genetics Reference Panel (DGRP), demonstrating the genotype-to-phenotype link in an independent set of diverse genetic backgrounds. The two SNPs lie in the putative regulatory regions of the genes *SP70* and *sog*, where allelic differences in gene expression correlate with the heat tolerance phenotype. Overall, our results suggest that regulatory loci that influence embryonic heat tolerance are under selection in nature. Our study extends previous work in developmental genetics of *Drosophila* by characterizing the genomics of an ecologically relevant developmental trait in natural populations.

**Significance Statement:** By comparing populations from distinct environments, we can determine the factors important for responses to environmental change. We used laboratory-based selection and whole-genome sequencing to uncover the genomic basis of embryonic heat tolerance in tropical and temperate fruit flies. We compared our results to published genomes of wild-collected flies from multiple continents and seasons. Strikingly, variants of two genes that influence embryonic heat tolerance are correlated with precipitation and temperature in natural populations. Further, different genetic variants produce distinct patterns of gene expression in response to heat stress. This is strong evidence that embryonic heat tolerance is under selection in nature and should be considered when forming predictions about responses to environmental change in species with complex life cycles.

## Introduction

For nearly a century, evolutionary biologists have studied how populations become differentiated despite the action of homogenizing forces like migration and purifying selection (*1–3*). This question has driven extensive research on local adaptation and spatially varying selection, providing strong evidence that selection structures natural populations (*4*, *5*). Species that span environmental gradients have been powerful model systems for the study of evolutionary and ecological genetics (*6–8*), in part because abiotic factors like temperature exert strong selective pressures that can be quantified (*9*, *10*) and, thus, the causative selective environments can be characterized.

The fruit fly, *Drosophila melanogaster*, is an excellent model for studying adaptation across ecological gradients. In addition to its well-established genomic tools and rich experimental literature, its natural populations possess characteristics that make them ideal for investigating the genetic basis of adaptation (*11*). For instance, *D. melanogaster* adults can overwinter in temperate regions, allowing for the establishment of resident populations that undergo local adaptation to their specific environments (*12–14*). This process contributes to the formation of adaptive clines along latitudinal and longitudinal gradients, many of which have been extensively studied (*6*, *15*, *16*). Among these, temperature clines are particularly well-documented (*17*), with notable examples in North America (*18*) and Australia (*19*).

Despite decades of research, uncovering the genetic basis of adaptation along temperature gradients remains challenging for several reasons. First, selection operates across both spatial and temporal scales. For example, *D. melanogaster* can rapidly adapt to seasonal changes through bursts of directional selection (i.e., adaptive tracking (*13*, *20–25*)). These adaptive dynamics are shaped by multiple, co-varying selection pressures across space and time, complicating the identification and experimental tracking of these changes (*14*, *26*, *27*). Second, in *Drosophilids*, genomic ancestry—rather than a shared selection history—primarily determines adaptive potential, so thermal tolerance genes identified in one population may not be relevant elsewhere (*28*). Lastly, selection acts throughout the organism’s life history(*29*). Indeed, our previous work has shown that thermal adaptation, particularly heat tolerance, is life-stage dependent and primarily occurs during the early embryonic stage, with little correlation to adult performance (*30*). This highlights the importance of assessing selection at the appropriate life stage to accurately measure its impact.

Given this complexity, identifying the agents of selection and their target genes requires integrating ecological and evolutionary insights with experimental validation. This approach is crucial for addressing several key questions: What are the primary agents of selection? At which stages of life history does selection act? And, most importantly, what are the causal loci responsible for thermal adaptation? In this paper, we tackle these questions by combining classical quantitative genetics methods with genomics, transcriptomics, and simulations to uncover the genetic basis of embryonic thermal tolerance. Specifically, we used an advanced introgression–backcrossing design (*31–33*) to map the genomic basis of embryonic heat tolerance in neotropical *D. melanogaster*, comparing lines from Vermont, USA (**VT**; temperate) and Saint Kitts, Caribbean (**SK**; tropical; **Fig. 1A**, **Table S1**). These two sites represent contrasting ecological regimes for flies (i.e., seasonal in VT and tropical in SK; **Fig. 1B–D**).

**Figure 1.**
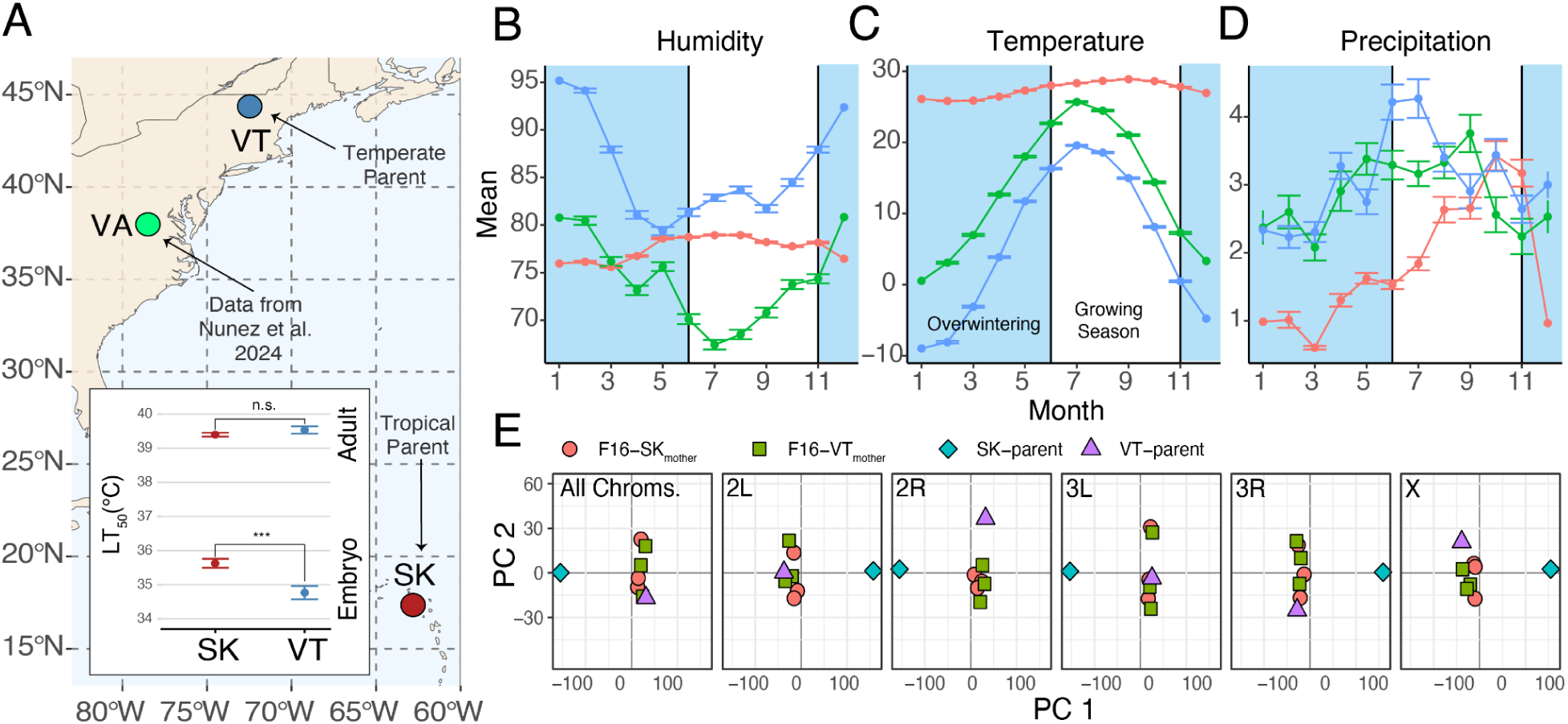
The ecological context of temperate and tropical *Drosophila*. **(A)** Map of North America showing the location of the parental tropical (SK) and temperate (VT) lines. Published samples from Virginia (VA) are also shown. **(A-inset)** LT_50_ of adults and embryos estimated from survival after acute heat stress, as reported in our previous study (***Wald test, *P* = 0.0003). **(B)** Average monthly humidity (%) over 25 years of data (2015–2025). The blue bands indicate the overwintering period of temperate *Drosophila*. **(C)** Same as B, but for mean air temperature (°C at 2 m). **(D)** Same as B, but for mean precipitation (mm/day). **(E)** Principal component analysis using parental and F16 lines. PCAs were estimated using all chromosomes, as well as for each chromosome separately. The percentage of variance explained (PVE) by PC 1, across analyses are 84.3% (All), 90.8% (2L), 85.9% (2R), 76.9% (3L), 85.4% (3R), 89.7% (X). The PVE by PC 2, across analyses are 4.22 (All), 3.33 (2L), 6.28 (2R), 8.18 (3L), 5.65 (3R), 3.17 (X).

Furthermore, these populations were used in a previous study showing that embryonic heat tolerance in SK is higher than in VT, yet adults exhibit comparable heat resistance, thus creating a natural experiment for dissecting the genetic basis of life stage-specific thermal adaptation (*30*). We integrated our results with data from the *Drosophila* Evolution over Space and Time (**DEST**) dataset, which encompasses samples from more than 520 populations worldwide (*14*, *34*). We also validated mutations of interest by measuring embryonic heat-shock survival in lines from the *Drosophila* Genomic Reference Panel (**DGRP** (*35*)), as well as quantified temperature-dependent gene expression.

## Results

### Advanced introgression maps embryonic heat tolerance to regions in 2R and X

We conducted a 16-generation selection and backcrossing experiment between VT and SK flies. We alternated even generations of heat-shock selection (∼80% mortality in ∼1-hour-old embryos) and backcrossing (to the VT genetic background) with odd generations of free recombination. This design is expected to produce a genome largely representative of the heat-sensitive VT parental population, except for tropical genomic regions associated with embryonic heat-shock tolerance. Indeed, after 16 generations of experimental evolution, the introgressed populations exhibited elevated embryonic heat tolerance (**Fig. S1**; Welch’s t-test, *P* = 0.013), whereas their genomic backgrounds, genotyped using Pool-Seq (*36*) (see **Fig. S2**), retained high similarity to the VT parent (**Fig. 1E**, see “All Chroms.”). Notably, this pattern is not observed in chromosomes 2R and X, where both SK/VT parents and the F16 pools cluster separately in PC space (**Fig. 1E, Fig. S3**).

To discover the genetic basis of embryonic heat tolerance, we measured genetic differentiation (***F*_ST_**) across pools and benchmarked it against population genetic simulations based on drift-only expectations (**Fig. 2A; Fig. S4A**). Overall, we observe 23 windows across the genome where all F16 pools outperform 95% of simulations (**Fig. S4A**) with the strongest signals detected on 2R and X (**Fig. 2A; Fig. S4B**). Indeed, this analysis revealed two windows of introgression that are consistently the top hits across all replicates: a 1.5 Mb window in 2R (19,115,753–20,615,753), near the right breakpoint of the cosmopolitan inversion *In(2R)NS*, and a 1 Mb region in X (15,123,410–16,123,410).

**Figure 2.**
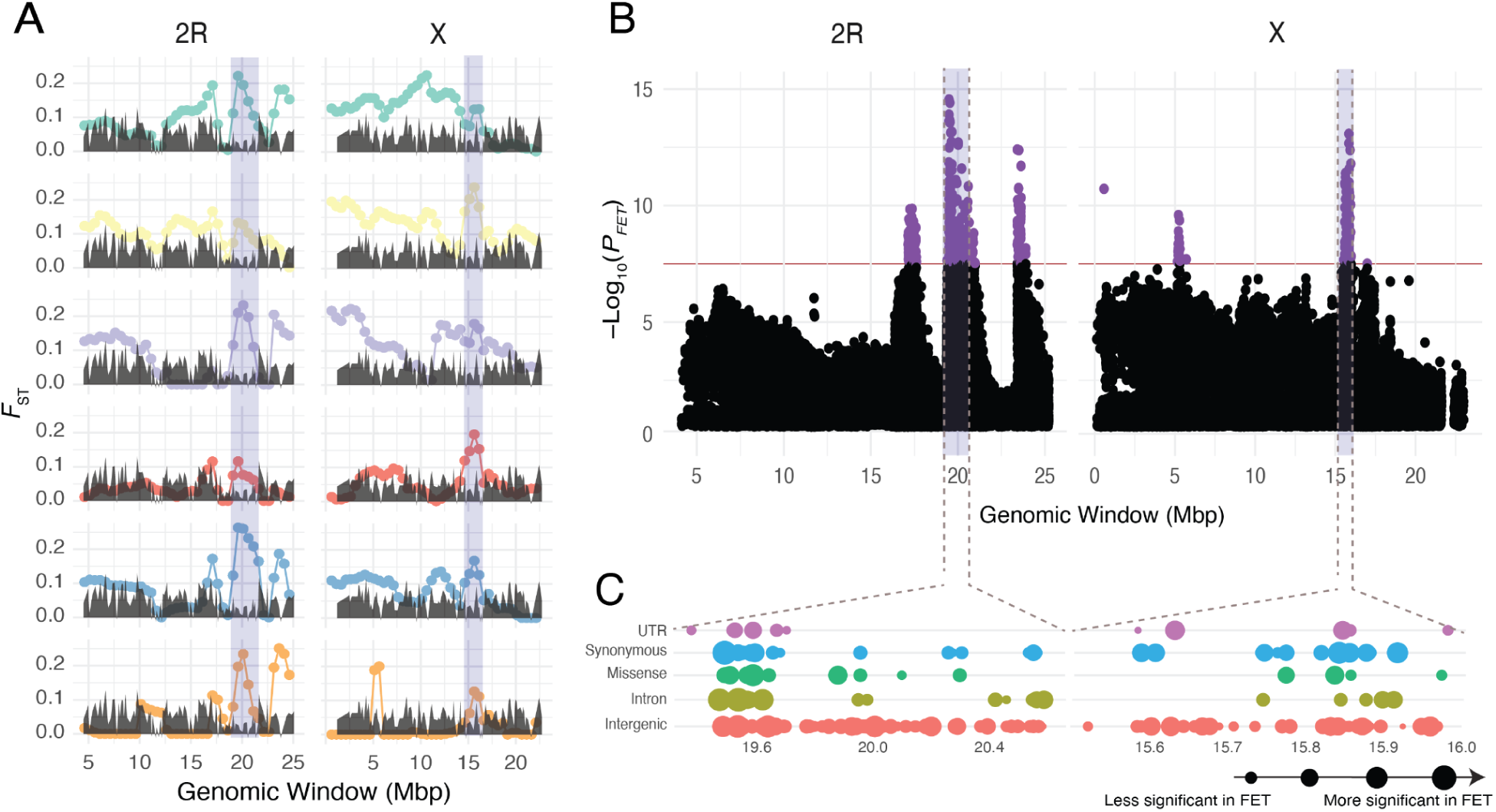
Genome scan for targets of embryonic thermal adaptation. **(A)** Sliding window analysis for *F*_ST_ outliers across our introgression crosses. The grey envelopes show the neutral introgression simulations. Significant regions are shown in purple. **(B)** Fisher’s exact test (FET) *P*-value (Bonferroni corrected) for the synthetic pool composed of all F16 replicates. The significant regions from panel A are shown in purple. The red line represents a threshold of ∼10^-8^ (equivalent to a Bonferroni cutoff for 0.01). **(C)** Functional annotations for the SNPs within windows of high concordance. The size of the circle indicates the FET *P*-value for the individual mutation.

To generate a list of putative targets of selection we applied a series of SNP-wise Fisher’s Exact tests (**FET**) on a “synthetic pool” created by aggregating all F16 samples, maximizing statistical power across replicates. This analysis identified 391 SNPs across the genome with significant FET *P*-values (0.01; Bonferroni corrected). Notably, the most significant FET hits are aligned with the top genomic windows identified in our *F*_ST_ analysis in 2R and X (purple bars in **Fig. 2B**).

The window of interest in 2R, at 19.5Mb, contains 157 genes and harbors 12,949 SNPs out of which 197 were associated with heat tolerance. Of these, 20 are missense mutations, 55 are synonymous variants, and 21 occur in UTRs. On the other hand, the window in X at 15.5Mb, contains 60 genes and harbors 2,673 SNPs, 58 that are associated with heat tolerance. Of these, 6 are missense mutations, 29 are synonymous variants, and 8 occur in UTRs (**Fig. 2C; Tables S2;** see **Text S1, Tables S3-S5** for additional information on functional annotations).

### Seasonal and latitudinal genome scans pinpoint embryonic heat tolerance SNPs

The strong concordance among our analyses highlights regions on 2R and X as the putative basis of embryonic heat tolerance, although the resolution remains coarse. To refine our mapping, we cross-referenced these candidate regions with published datasets to identify SNPs previously reported as clinal (*21*) or seasonal (*23*). Because most ecological genetics studies focus primarily on autosomal variation, we first limited our co-localization analysis to SNPs on 2R. We identified 11 SNPs associated with both clinality and embryonic thermal tolerance (**Fig. 3A**) and two SNPs linked to seasonality (**Fig. 3B**). Strikingly, one SNP, 2R: 20,551,633 (C→A on the forward strand; G→T on the reverse strand), is significant in both clinal and seasonal studies (**Fig. 3C**) as well as in our embryonic heat tolerance analyses.

**Figure 3.**
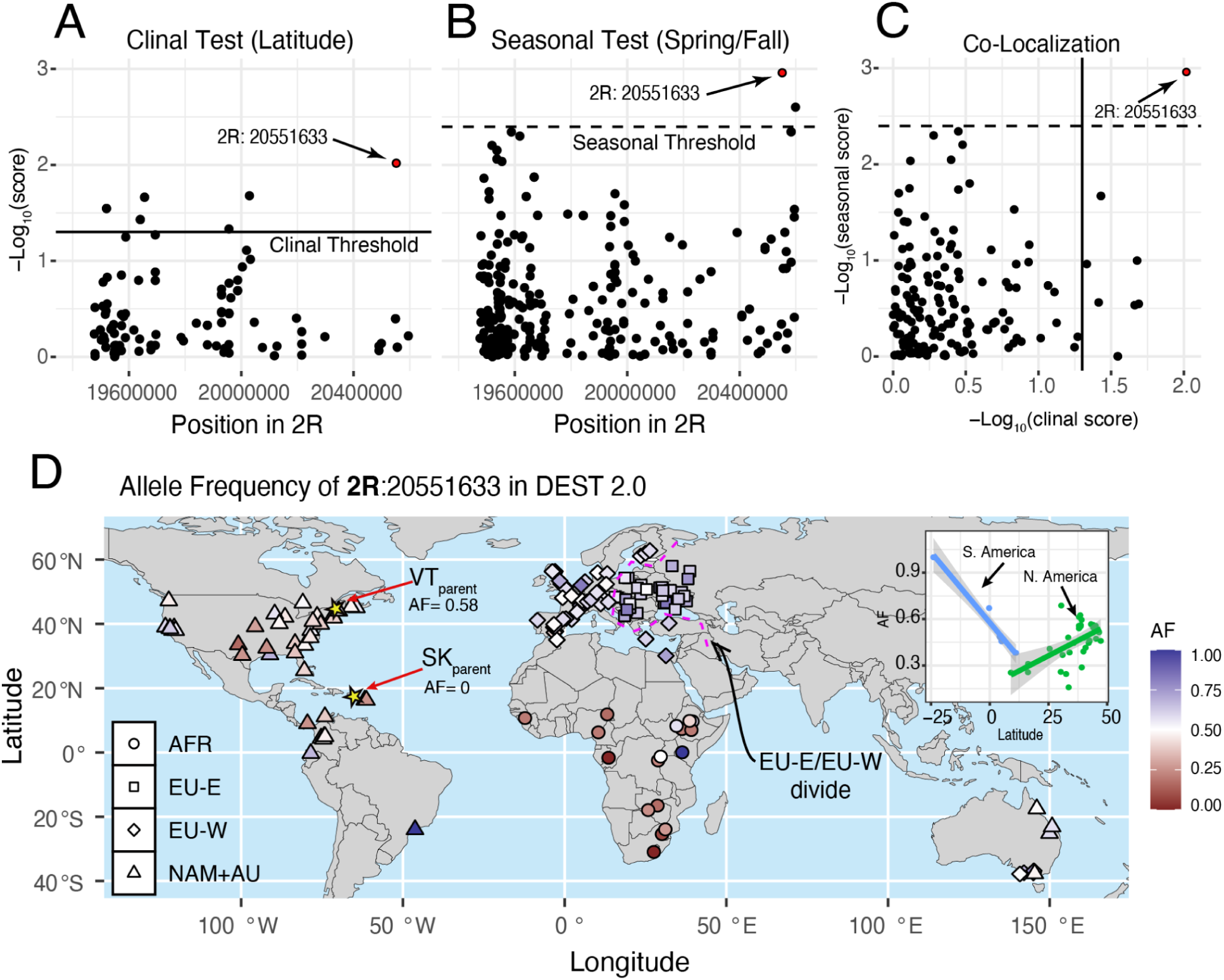
Footprints of seasonality and clinality at the window of interest in 2R. **(A)** Colocalization analysis of outlier SNPs in the FET analysis with clinal SNPs, the horizontal line is the significance threshold indicated by the authors. **(B)** Colocalization analysis of outlier SNPs in the FET analysis with seasonal SNPs, the horizontal line is the significance threshold indicated by the authors. **(C)** SNPs that are jointly FET outliers, seasonal outliers, and clinal outliers. **(D)** Global allele frequencies at the top SNP in 2R (position 20,551,633; frequency of A shown) from the DEST 2.0 dataset. Shapes indicate known demographic clusters in *D. melanogaster* (AFR: Africa; EU-W: Europe West, EU-E: Europe East; NAM+AU: North America and Australia)**. (D-inset)** Patterns of allele clinality in North (green) and South (blue) America.

In our experimental lines, we observed that the SNP is fixed for the “C” allele in the SK parent (i.e., 2R^C^ is tropical) and that it is polymorphic in the VT parent; yet, A is the major allele (AF_C-allele_ = 0.598; i.e., 2R^A^ is temperate). Whereas the F16s pools show a mean allele frequency of the A allele of 0.18. This allele frequency is substantially lower than expected (0.021%) under the VT introgression design (*p*_expected_ = 0.59; Δ*p* = 0.41; see p. 406 of (*32*)). This large shift (41.7%) is strong evidence that this SNP played a major role in embryonic heat tolerance during adaptive introgression.

To assess the potential ecological relevance of this SNP, we explored its patterns of variation in the DEST 2.0 dataset (*14*). We detected strong clinal patterns across the Americas (**Fig. 3D**), with allele frequencies increasing with latitude in North America (*r* = 0.540, *P* = 0.0012; **Fig 3D-inset**) and decreasing in South America (*r* = -0.9816, *P* = 8.70×10^-5^). We also performed a locus-specific test for seasonality, using temporal data from Charlottesville, Virginia (USA; **Fig 1A**, green dot) from a previously published study(*13*). Our results indicate that our mutation in 2R experiences seasonal selection associated with the variance of temperature, 0-45 days prior to collection (**Fig. 4A**; T_SD:0-45_; *β_seasonal_* = -0.089, *P_seasonal_* = 0.0053; beats 96% of permutations).

**Figure 4.**
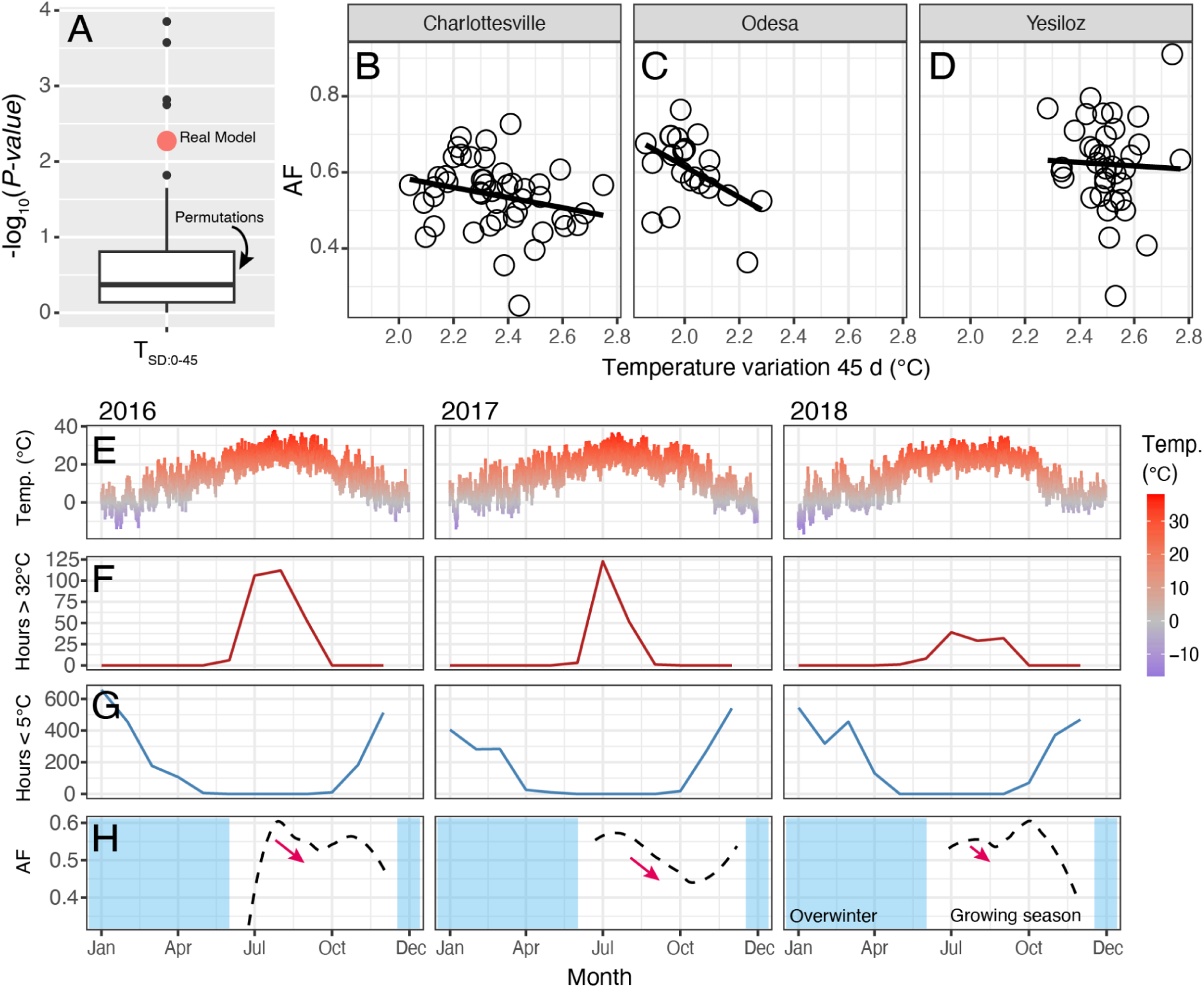
Dynamics of seasonal selection at the top SNP in 2R. **(A)** Result of the generalized linear model at the top SNP in 2R using genomic data from Virginia. The real estimate for the locus is shown in red, and the outcome of 100 permutations is captured by the boxplot. The data shown represents the model summarizing temperature variance 0-45 days prior to collection. **(B)** Allele frequency relative to temperature variance 0-45d in Charlottesville, Virginia. **(C)** Same as B, but in Odesa, Ukraine. **(D)** Same as B, but in Yesiloz, Turkey. **(E)** Daily mean temperature in Charlottesville, Virginia from 2016-2018. **(F)** Number of hours per month above 32°C. This is a proxy for summer-like conditions. **(G)** Number of hours per month below 5°C. This is a proxy for winter-like conditions. **(H)** Allele frequency trajectories of the A allele across the growing seasons 2016-2018. The overwintering period is shown in blue.

The timeframe of 0-45 days prior to collection suggests that populations with low A allele frequency likely descend from flies exposed to high temperature variability at least three generations earlier, when selection favored the C allele. To illustrate these dynamics, we plotted allele frequency versus T_SD:0-45_ using the seasonal data from Virginia (**Figs. 1B-D**) and extended this analysis to two additional DEST populations with high-resolution seasonal sampling: Odesa, Ukraine and Yesiloz, Turkey. Our results show that allele frequencies vary in relationship to T_SD:0-45_ in Charlottesville (*r* = -0.2985, *P* = 0.072; **Fig 4B**), and Odesa (*r* = -0.4627, *P* = 0.026; **Fig 4C**), but not in Yesiloz (*r* = -0.038, *P* = 0.82; **Fig 4D**). Two insights are derived by plotting weather data (**Fig. 4E**) relative to the allele frequencies from Virginia (**Fig. 4H**). First, allele frequencies shift after late-fall cold snaps (T<5°C; **Fig. 4G**), but their year-to-year direction is inconsistent, likely driven by drift during winter population bottlenecks(*13*). Second, the SNP shows a consistent pattern of summer selection whereby the A allele decreases in frequency (**Fig. 4H**; **Fig. S5**) following heat stress (i.e., days T > 32°C, as per the thermal limit model (*23*); see **Fig. 4F**). Importantly, this direction of allele frequency change mirrors what we saw in our adaptive introgression experiment. Overall, these data suggest that 2R:20,551,633 (or a closely linked variant) is likely an ecologically important mutation.

Repeating our analyses on the X chromosome revealed six loci with significant signals in both seasonal and clinal tests (**Fig. S6A**). Evaluating the pattern of introgression in our F16 pools revealed that three of these six loci showed clear patterns of adaptive introgression (positions: 15,602,941; 15,607,604; and 15,847,814; **Fig. S7**). Of these three loci, mutation X:15,607,604 emerges as a candidate locus showing a signature of seasonal evolution (*β_seasonal_* = -0.061, *P_seasonal_* = 0.045 [beats 90% of permutations]) relative to maximum precipitation (0-45 days prior to collection). The candidate locus in X has two alleles (A→T; see **Text S2** for more details), with “T” fixed in SK (i.e., X^T^ is tropical) and present at low frequencies (∼0.05) in VT (i.e., X^A^ is temperate). In the introgression experiment, the locus increased to an average of 0.38 (shift due to selection: Δ*p* = 0.33). Overall, while the association with thermal selection among natural populations in the DEST dataset is less evident for X:15,607,604, the observed signals of seasonal and clinal variation at this locus, paralleling that of 2R:20,551,633, suggests it also has ecological relevance.

### Linking allelic variation to embryonic survival in the DGRP panel

Thus far, we have identified two focal SNPs on 2R^C/A^ and X^T/A^ that align with top introgression hits and show strong environmental associations. To assess their direct links to embryonic fitness traits, we conducted a heat shock survival assay (45 mins. at 35°C) on one-hour-old embryos from 64 lines from the DGRP (**Table S6**). Since these alleles are not linked (*r*^2^_X-2R SNPs_ = 0.0008), all four possible homozygous combinations were tested (i.e., one allele from each climate, 2R^C^X^A^ = 22 and 2R^A^X^T^ = 13; both tropical, 2R^C^X^T^ = 8; and both temperate: 2R^A^X^A^ = 21). Overall, we observe high levels of variation for this trait in the DGRP, with line survival ranging from 0% to 66%, showing that the trait is polygenic (**Fig. 5A**). Yet, consistent with our introgression results, genotypes homozygous for tropical alleles (2R^C^X^T^) showed significantly higher survival (**Fig. 5B**), relative to all other combinations (ANOVA; 2R^C/A^-by-X^T/A^ effect, *F*_1,55_ = 11.22, *P* = 0.0014; *Wolbachia* and inversions [*In(2L)t* and *In(2R)NS*] are not significant). Considered individually, X^T/A^ shows a marginal effect (*P* = 0.076; **Fig. 5D**), whereas 2R^C/A^ is nonsignificant (*P* = 0.162; **Fig. 5C**). These findings suggest that both tropical mutations (2R^C^ and X^T^) are required to enhance embryonic survival under heat stress, with their joint large effect indicating likely epistatic interactions (Note that this effect is not observed with the other two candidate mutations in X; **Fig. S6B-G**). We also used data from Lecheta et al. (2020)(*37*) to evaluate adult thermal performance phenotypes (CT_max_) in these DGRP lines. Strikingly, 2R^C^X^T^ lines showed a minor decline in thermal performance relative to other genotypes (**Fig. 5C-G**).

**Figure 5.**
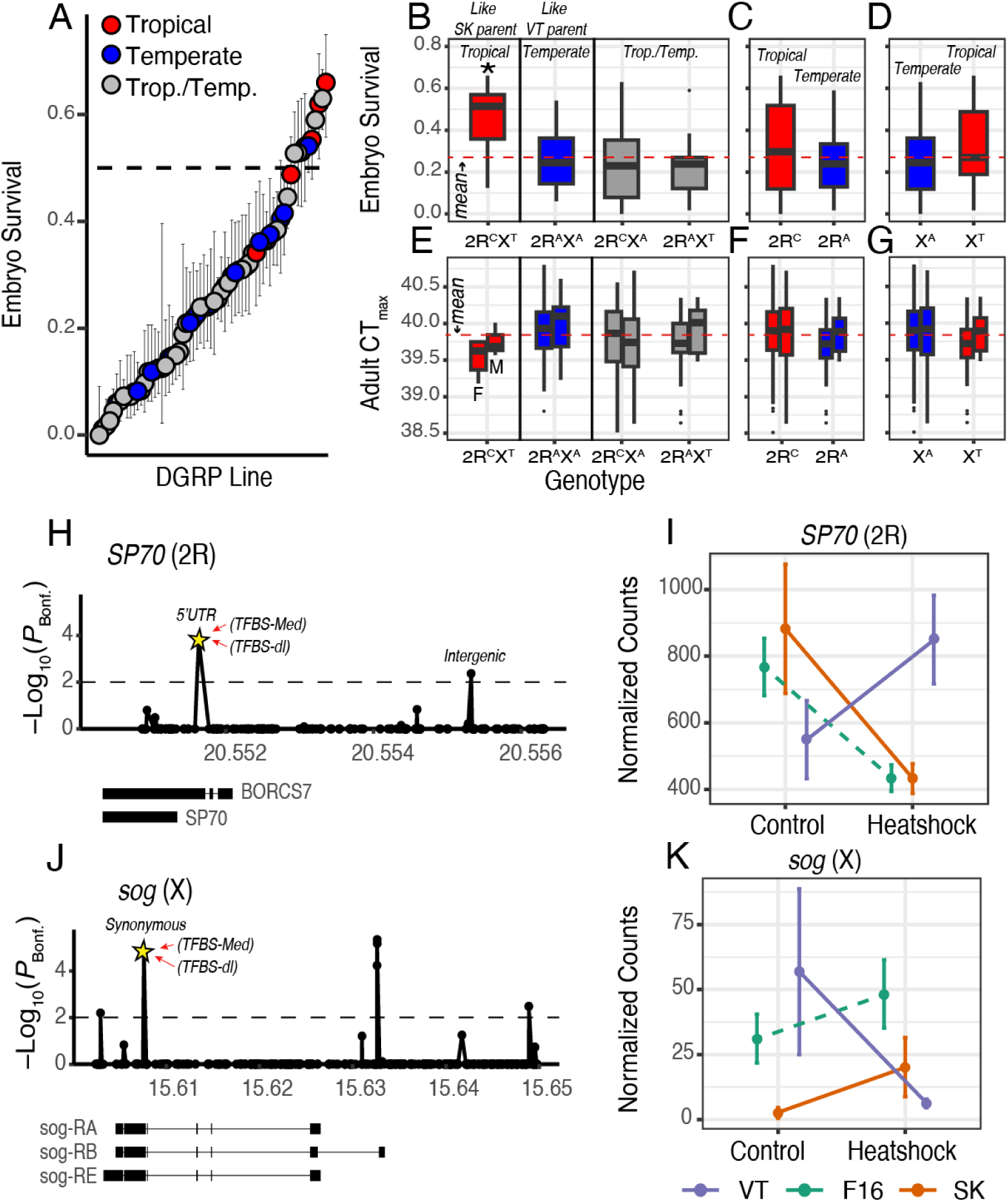
Genes and phenotypes associated with the top SNPs in 2R and X. **(A)** Embryonic survival (proportion surviving) for 64 DGRP lines. The colors indicate whether the lines are either the tropical or temperate homozygous of 2R^C/A^ and X^T/A^. Lines with one allele of each kind are shown in grey. For each line the 95% binomial confidence interval is shown. The horizontal dashed line represent 50%. **(B)** Embryonic survival for all genotypic combinations of 2R^C/A^ and X^T/A^. **(C)** Same as B but for only 2R^C/A^. **(D)** Same as B but for only X^T/A^. **(E-G)** Same as B–D, but showing adult thermal performance. In each plot, female measurements are on the left and male measurements on the right. **(H)** SNPs in *SP70* that are significant in the introgression FET analysis. The top seasonal and clinal SNP is indicated with a star. Feature annotations are also shown. **(I)** Levels of transcript expression (normalized read counts) for SK, VT, and F16 embryos, at 25°C (Control) or after a heat shock of 34°C for the *SP70* gene. **(J)** Same as H, but for *sog*. See Table S7 for feature annotations of regions in (H) and (J). **(K)** Same as I for the *sog* gene.

### Top embryonic SNPs map to genes interacting with *Medea* and *Dorsal*

To identify genes associated with our focal mutations (2R^C/A^ and X^T/A^), we integrated published developmental transcriptomes (*38*), genomic feature libraries (*39*, *40*), including published ChiP-seq data (*41*), and heat-shock RNA-seq generated for this study. Notably, 2R^C/A^ falls within a transcription factor binding hotspot with five transcription factor binding motifs (**TFBMs**), including the motifs of Medea *(Med)* and dorsal (*dl*), transcription factors involved in dorso-ventral patterning in the early stage embryo (*42*). Only considering annotations of genes with detectable expression levels at 0-2 hours post-egg laying (**Table S7**), the mutation is associated with two protein coding genes (**Fig. 5H**): *SP70* (CG13430; as a 5’UTR) and *BORCS7* (CG18065; as a synonymous variant; **Table S8**). Likewise, the top hit on the X chromosome is a synonymous variant in *sog* (short gastrulation; CG9224; **Fig. 5J**). Although the SNP does not alter the amino acid sequence of *sog*, it falls within another transcription factor binding hotspot with binding motifs for eight transcription factors, including *Med* and *dl*. Remarkably, both 2R^C/A^ and X^T/A^ occur within TFBMs for the same key developmental genes (i.e., *Med* and *dl*; **Table S8**).

To validate our findings, we generated an independent gene expression dataset from heat-shocked embryos of the same populations used in the introgression experiment (VT, SK, and the F16 offspring). Our findings show that *SP70*, but not *BORCS7,* was highly expressed and showed contrasting heat-shock responses in the VT and SK parental lines, with the F16s following the same reaction norm as the SK parent (**Fig. 5I**). Notably, *sog* expression mirrored that of *SP70* with F16 offspring showing the same expression as SK parents (**Fig. 5K**). Based on this collective evidence, we propose that *SP70* (2R^C/A^), predicted to encode a serine-type endopeptidase, and *sog* (X^T/A^), which encodes a secreted developmental morphogen, house key, large effect loci, underlying the genetic basis of embryonic heat tolerance in the wild.

## Discussion

### Mapping seasonal and clinal mutations to embryonic survival

The *SP70* and *sog* loci are notable for several reasons. Beyond the strong signal detected in our mapping experiment (**Fig. 2**), colocalization with DEST indicates that mutations in these two genes exhibit genomic footprints of both clinal (*6*), and seasonal (*27*) adaptation. These findings offer three insights into the genetic basis of adaptation in nature.

First, our data revealed significant clinality across multiple spatial scales. Yet, these patterns must be considered in the context of *Drosophila*’s phylogeography, as North American populations arose from a secondary contact between old world flies (*14*, *43*), which could generate neutral clines. However, our experiments show that these mutations enhance embryo survival, suggesting that the clines may be adaptive. This aligns with previous studies that exemplify adaptive clines along the east coasts of North America and Australia (*44*, *45*).

Second, beyond strong latitudinal clines, our genomic mapping provides evidence for two loci undergoing seasonal adaptive tracking(*20*). While adaptive tracking affects thousands of *Drosophila* genes, few loci have been experimentally validated or linked to ecologically important phenotypes (*13*, *21*, *23–25*, *46–48*) (but see (*26*, *49–52*)). In this context, our findings provide empirical evidence for adaptive loci linked to embryonic survival. For example, in our introgression, *SP70* shifted from the temperate allele (2R^A^) to the tropical allele (2R^C^) after 16 alternating generations of heat selection. This pattern was also seen in DEST (e.g., Virginia, Ukraine, Turkey), where the 2R^A^ allele tracks temperature changes 1.5 months prior to collection (**Fig. 4A, H**).

Third, SP70 and sog alleles are remarkable in their consistent responses to environmental variables across space and time. This is remarkable given that only ∼3.7% of genomic loci exhibit parallel seasonal and clinal changes in previous studies (*53*). In a landscape where finding loci with consistent ecological responses is akin to finding a needle in a haystack, the ability to pinpoint such targets represents a significant step forward in our understanding of the genetic basis of adaptation.

### Selection acts differently across life stages. What does this mean for polygenic adaptation?

Beyond revealing spatial and temporal selection patterns, our work suggests that the embryo is a crucial stage for adaptation, as embryos are immobile and particularly vulnerable to environmental stressors (*54*, *55*). Differences in mobility between early and late life stages can cause the same stressor to produce distinct selection dynamics across development. Indeed, these dynamics can lead to complex patterns of selection, such as ontogenetic decoupling (*56*) or antagonistic pleiotropy (*57*, *58*), both processes that have been documented across various taxa: insects (*29*, *59–61*), amphibians(*62*, *63*), birds and mammals (*64*, *65*), and marine invertebrates (*55*, *66*). Indeed, our data show that although genetic variation for embryonic heat tolerance is ample, the tropical alleles at *SP70* and *sog* enhance embryonic heat tolerance early in development, while reducing thermal performance in adults, consistent with antagonistic pleiotropy. Moreover, neither allele alone improves survival, but together they produce large effects, indicating epistasis.

We note that previous quantitative genetic studies of thermal tolerance have demonstrated that the trait is highly polygenic (*37*, *67*, *68*); yet, here we identify two alleles of large effect. How can this be reconciled? One explanation is that the introgression mapping experiment, while representing genetic variation across a broad geographic region, only consisted of two genetic backgrounds, and thus we present herein one part of what may be a more complex story underlying the genetics and evolution of embryonic thermal tolerance. On the other hand, our introgression mapping results are consistent with a polygenic basis of embryonic heat tolerance, as 23 regions of the genome significantly contributed to variation in the trait in at least one replicate introgression. Meanwhile, the two loci of largest effect were the genomic regions that were consistently targets of selection across all six replicates, suggesting that while embryonic heat tolerance has a polygenic basis, it may differ from adult heat tolerance in the relative contribution of large- vs. small-effect loci (*37*, *67*, *68*). Moreover, epistatic effects may be missed in GWAS (*69*), as each locus in isolation produces no measurable effect on the phenotype. Thus, it could be that large-effect loci underlying thermal traits in later life stages remain to be discovered. Further clues may come from the functions of the target genes *SP70* and *sog*, which we outline below, suggesting a physiological difference between embryonic and adult heat tolerance that may influence the genetic architecture that underlies each trait. Overall, the data presented herein support our previous work (*30*) and suggest that early life stages play a key role in the ecological physiology of species with complex life cycles.

### Why are embryonic and adult thermal tolerances decoupled?

Differences in life-stage responses to the same selection likely arise because adults, unlike embryos, possess distinct mechanisms to cope with stress, such as behavioral thermoregulation (*70*). Consequently, thermal selection in adults often acts on traits beyond survival. For instance, traits like male fertility (*71*) correlate more strongly with environmental temperature than adult heat tolerance. There is even evidence that selection acts on the circadian rhythm of egg laying in *D. melanogaster*, such that populations that experience extreme daytime heat only lay eggs at night (*72*). To our knowledge, there is no evidence of selection on female choice of oviposition site in egg laying behavior (*73*). Further, the thermal properties of necrotic fruit (*74*) and *in situ* temperature data (*9*) suggest that the thermal microenvironment of the *Drosophila* embryo is highly variable and likely to exceed the temperatures that cause thermal stress on a regular basis (*30*, *75*, *76*). Thus, because embryos are immobile, they are likely to be at the mercy of the microenvironment in which they are laid. Accordingly, we suggest that embryonic thermal traits are a heretofore underappreciated aspect of thermal adaptation. In fact, it is plausible, in the context of genetic correlations, that adult thermal traits track environmental change as correlated responses to selection on embryonic traits, and this may be the reason for the comparatively weak correlation of adult heat tolerance and environmental temperature (*30*, *77*). Furthermore, if thermal traits are genetically and/or physiologically decoupled across life stages, then the most thermally sensitive life stages could be key to ultimately establish the upper thermal limits of a species more broadly (*78*).

### Gastrulation as a putative physiological target of selection

Our data, in the context of *Drosophila* embryogenesis, may reveal molecular targets of thermal selection. We assessed heat tolerance in embryos under 2 hours old, when the body axis is established and primordial cell types begin specifying tissues and organs, including gastrulation, a complex process regulated by interacting genetic factors (*79*, *80*) that orchestrate cell shape changes, cell migration, and tissue and organ formation (*81*, *82*). Disruptions to gastrulation lead to developmental arrest (*80*). Importantly, gastrulation takes place within an hour of the developmental window in which we characterized embryonic heat tolerance (*83*). Based on this developmental context, the functions of our two major effect loci (*SP70* and *sog*), and how these genes were expressed in heat-tolerant vs. heat-sensitive embryos, we hypothesize that regulation of gastrulation is the physiological target of thermal selection.

Previous work suggests that *SP70* and *sog* interact indirectly to regulate gastrulation (*84–86*), with *SP70* promoting gastrulation and *sog* inhibiting it. Although the function of the SP70 protein has not been characterized in *D. melanogaster*, the human ortholog, TMPRSS4, induces the epithelial-to-mesenchymal transition (EMT)(*87*), which is a pro-gastrulation process that facilitates mesoderm formation via cell specification and migration (*86*). The interaction between *SP70* and *sog* is highlighted by *sog*’s direct inhibition of the secreted factor *Dpp*, which in turn activates the pMad/Medea transcription factor complex (*42*, *85*). Strikingly, both our top SNPs in *SP70* and *sog* lie in the binding motif of pMad/Medea (*41*, *88*) suggesting that these two genes are co-regulated. These findings, along with our RNA expression data, suggest that *sog* inhibits the expression of *SP70*, a pro-gastrulation signal.

Recall that *sog* and *SP70* showed anti-correlated expression: heat-tolerant embryos increased *sog* and decreased *SP70* after heat stress, suggesting they slow or inhibit gastrulation to recover from stress. In contrast, heat-sensitive embryos decreased *sog* and increased *SP70*, likely promoting gastrulation during heat shock. Proceeding through gastrulation during heat stress could be disastrous due to (i) thermally induced disruptions to cell membranes (*89*) and proteins (*90*) that orchestrate gastrulation and (ii) misexpression of genes involved in the spatial control of gastrulation, potentially causing organ malformations (*91*, *92*). Thus, adaptive embryonic heat tolerance may involve the halting of gastrulation, akin to developmental quiescence, that serves to protect vulnerable morphogenetic processes. Such a mechanism may be distinct from thermal adaptation of acute heat tolerance in adults, which involves the regulation of many genes, including molecular chaperones (*93*) and genes of the nervous system (*37*, *67*, *68*).

Why, then, do the alleles that confer the pro-gastrulation response under heat shock conditions segregate in nature? We hypothesize that more benign increases in temperature, if they don’t lead to thermal denaturation of lipid membranes and proteins, could create ideal conditions for gastrulation to occur and thereby speed up development time (*94*). Shorter development time may be advantageous because it limits the amount of time that flies exist in a vulnerable life stage (*54*), as well as shortening the generation time. While it is beyond the scope of the present study to test our molecular physiological model of embryonic heat tolerance mediated by gastrulation signals, future work will capitalize on this unique opportunity to uncover the mechanism of adaptation to environmental variability in embryos.

In conclusion, by integrating quantitative genetics and ecological genomics, we provide, to our knowledge, the first genomic mapping of an ecologically relevant trait in *Drosophila* embryos. Loci showing both spatial and temporal adaptation are rare, yet we identify two—*SP70* and *sog*—on separate chromosomes, highlighting the ecological relevance of embryonic heat tolerance. Gene expression data, together with previous work, suggest these genes interact to coordinate early development under environmental variability. Furthermore, while most studies of *Drosophila* heat tolerance focus on adults, few link genotype to phenotype in an adaptive context; our results fill this gap and offer a novel perspective on ecological and evolutionary physiology across the *Drosophila* life cycle.

## Methods

### Fly lines

As we previously reported, embryos (but not adults) of several tropical lines, collected from a broad geographic span around the world, have higher heat tolerance than lines collected from three populations in Eastern North America (*30*). In order to map the genomic basis of variation in embryonic heat tolerance, we first screened these tropical and temperate North American lines for compatible inversion genotypes among the 9 major cosmopolitan inversions known to segregate in *D. melanogaster* populations (*95*). We chose two parental lines with compatible inversions, one tropical line from the island of Saint Kitts (SK) in the Caribbean and one temperate line from Vermont (VT), USA. These two lines possess the standard inversion genotype at all positions, except for In(3R)Payne where they possess the inverted genotype. We originally obtained the Vermont parental line as a gift from KL Montooth (Stock name: VTECK8). This line was established from a wild collection of a single female from East Calais, VT (44.4°N, -72.4°E) and was subsequently isogenized by full-sib inbreeding for several generations (*96*). We obtained the tropical parental line from the *Drosophila* Species Stock Center (Stock: 14021-0231.34). This line was established from a wild collection of a single female from Monkey Hill on the island of Saint Kitts (17.3°N, -62.7 °E). We maintained flies under common-garden conditions fed with cornmeal-yeast-molasses medium at 25°C and 12L:12D for several generations prior to introgression mapping.

### Environmental data

We obtained hourly estimates for three environmental parameters, temperature, humidity, and precipitation, from the NASA POWER dataset (*97*). The data covered the period from January 1, 2015, to January 1, 2025, and were downloaded for three locations: sites corresponding to our parental lines (VT and SK) and a site in Virginia (VA) from (*13*). Specifically, we retrieved data for VT (latitude: 44.36272, longitude: -72.47167), SK (latitude: 17.37319, longitude: -62.80985), and VA (latitude: 37.97900, longitude: -78.48970).

The NASA POWER dataset provides meteorological variables at a spatial resolution of 0.5° × 0.625° of latitude and longitude, respectively. We queried relative humidity (RH2M), which represents the ratio of actual water vapor pressure to the saturation vapor pressure at 2 meters above the surface, expressed as a percentage. For temperature, we used T2M, which estimates the average air (dry bulb) temperature at 2 meters above the surface. Lastly, precipitation was obtained using the PRECTOTCORR variable, which provides a bias-corrected estimate of total precipitation at the surface, including the water content in snow, expressed in millimeters per day.

### Introgression mapping

Embryos of SK are more heat tolerant than embryos of VT, with LT_50_s of 35.63°C and 34.77°C, respectively, following an acute exposure to 45 min of heat stress (*30*). While this difference in LT_50_ is approx. 1°C, this represents a significant difference in survival following heat stress. For example, a heat stress of 36°C induced 87% mortality in the VT line but only 63% mortality in the SK line (*30*). Tolerance to sudden acute increases in temperature is likely to be an ecologically relevant trait for embryos developing on necrotic fruit, where rates of temperature increase can exceed +15°C h^-1^ (*98*). To map the genomic basis of this variation in embryonic heat tolerance, we conducted an advanced introgression with selection and backcrossing design (*31–33*). We replicated the introgression 6 times, and we started each replicate introgression by crossing 300 SK flies with 300 VT flies. We did reciprocal crosses, such that three replicate crosses used SK founder females and the other three replicate crosses used VT founder females. We included this reciprocal mating design to account for the effects of the maternal cytoplasm (e.g., mitochondria) on embryonic heat tolerance. F1 hybrids were maintained at a population size of 1500 and allowed to mate freely, thus allowing free recombination. F2 progeny were collected on grape juice agar plates at 25°C at 0-1 h post-fertilization and immediately exposed to an acute heat shock. We followed the protocol of (*30*) except that embryos were heat-shocked for 1 hour instead of 45 min. The heat shock temperature was 35.2±0.05°C (mean±s.d.), which we reassessed prior to every selection bout to induce 80% mortality (20±10% survival). Survival was scored after 24 hours at 25°C and survivors were reared to adulthood. 300 virgin female survivors were then backcrossed with males of the heat-sensitive VT, and the progeny were allowed to mate freely. We carried out this scheme of free recombination every odd generation and selection and backcrossing every even generation through F16. At generation 16, we again heat-shocked embryos and collected 30 adult females from the survivors for pooled whole-genome sequencing.

### DNA sequencing, read mapping and SNP calling

DNA was extracted with a phenol-chloroform extraction protocol (*99*). Each introgression replicate was sequenced on the Illumina NovaSeq 6000 using a configuration of 2×150 (insert size 350). DNA reads were mapped to a hologenome for *D. melanogaster* (i.e., the genomes of *D. melanogaster* and known commensals and parasites combined) using the self-contained mapping pipeline developed for the DEST dataset (*34*). Mapping statistics for each pool were evaluated using qualimap v2.2.1 (*100*). SNP calling was done using PoolSNP (*101*), as part of the DEST calling pipeline. We ran PoolSNP with the following parameters: minimum allele frequency (maf) = 0.001, minimum allele count (mac) = 5, minimum coverage (min-cov) = 4, maximum coverage (mac-cov) = 95% (the dataset’s overall coverage), missing fraction (miss-frac) = 0.5. The resulting variant call file (VCF) was annotated using SNPeff’s *Drosophila melanogaster* library “BDGP6.32.105.”

### Estimation of genetic diversity, PCA, and introgression analyses

Analyses of nucleotide diversity (π) were done using npstat v1.0 (*102*). Calculations were done directly from the bam files outputted from the DEST pipeline and processed using samtools v1.10 (*103*). We used principal component analysis (PCA) to assess the results of the introgression process. To assess introgression at a global level, we used FactoMineR (*104*) to calculate PCA on the genomic data including the two parental pools and all six of the F16 pools. Then, to assess admixture across the genome, we repeated the PCA process using a sliding window scheme (window size = 0.1 Mb; step = 50 Kb). We quantified the genomic background of each F16 line as either more “VT-like” or “SK-like” by calculating the ratio between the mean euclidean distance (i.e., *d*), in PCs 1 and 2, of each F16 pool relative to the SK or VT parent. In this analysis, the Log_2_(*d*_SK_/*d*_VT_) value is a proxy for the genetic distance between the F16s and the parents, whereby Log_2_(*d*_SK_/*d*_VT_) > 0 indicates higher similarity to the VT parent and Log_2_(*d*_SK_/*d*_VT_) < 0 indicates higher similarity to the SK parent.

### Simulating neutral introgression in a non-Wright-Fisher framework

We used computer simulations to assess whether the observed patterns of genetic variation in our F16 crosses differed from simple expectations of genetic drift. This model assumed that no introgression took place and thus that all patterns of genetic variation resulted from genetic drift. To this end, we used the forward-in-time simulator SLiM [v4.2.2] (*105*), and a non-Wright-Fisher framework (*106*), to simulate 16 generations of drift with rounds of back-crossing into a parental line (similar to the real experimental design). Accordingly, we simulated two populations (*j*_1_ and *j*_2_) of size ∼1,500 each. In SLiM, non-Wright-Fisher models operate using carry capacities and thus the real census sizes of the population vary around 1500. We simulated 608 windows as “virtual chromosomes” emulating the real genome of *D. melanogaster* with length and mean recombination rate (ρ) equal to that of the real window. In the case of population *j*_1_, we created mutations based on the allele frequency vector of the synthetic cross, *p*_iF1_, for population *j*_2_, mutations were populated only based on a vector of the VT parental frequencies (i.e., *p*_i-VT_). The first step of the simulation is to generate a synthetic cross between two virtual “VT” and “SK” lines. Based on our crossing design, we assume that the per locus allele frequencies (*p*_i_) of the F1 cross can be modeled as the mean allele frequency of each parental line (eq. 1):

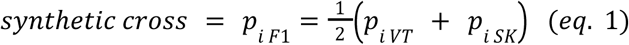

where “*p_i-K_* “ is the frequency of allele “*i*” in population “*K*” (i.e., either F_1_, VT, or SK). In this case, the initial allele frequencies of VT and SK are derived from the sequenced parental pools (i.e., SK_0_1 and VT8_0_1). Next, we divided the genome into regions with similar recombination levels using the data from (*107*). In total, we divided the genome into 608 windows (110 in 2L, 128 in 2R, 110 in 3L, 137 in 3R, and 123 in X). On average, these windows spanned 0.15 Mb (s.d. = 0.21 Mb). Accordingly, we simulated each of the 608 windows as a “virtual chromosome” with length and mean recombination rate (ρ) equal to that of the real windows. We also applied a mutation rate (μ) of 1.0×10^-7^, yet this parameter is not impactful in the simulations due to the small number of generations. In this model, the events of odd generations are unchanged. Yet, at even generations, in addition to the 80% bottleneck, the population experiences an introgression event (*j*_2_ → *j*_1_) where 300 individuals with *p*_i-VT_ are introgressed into the surviving population. While in the drift-only model, the bottleneck occurs via an instantaneous population size change, in this model, the contraction occurs by randomly choosing 80% of individuals to die in a *first()* callback event. Subsequently, during the same *first()* callback event, 300 individuals from *j*_2_ are moved into *j*_1_ and allowed to reproduce back to full carry capacity. This cycle repeats itself for 16 generations. We simulated 100 replicates for each recombination window across the genome for a total of 60,800 “neutral-introgression” simulations.

We also simulated a final step where genotypes are measured by applying pool-seq-like noise to the simulation’s outcome. To this end, we applied a two-step binomial sampling to the output of the simulations [see (*108*)]. In the first binomial step, we created a vector of allele frequencies equivalent to sampling 30 diploid individuals (i.e., 60 chromosomes) from the simulations. In the second binomial step, we simulated the sequencing effort by creating a vector of allele counts using a coverage estimate of 85X (similar to the real experiment). Notice that the allele frequency used in the second step is the output of the first step.

### Identifying regions enriched for large F_ST_ outliers

We assessed signals of genetic differentiation using the fixation index (*F*_ST_) statistic, as defined for pool-seq in Hubert et al. (*109*), implemented in the *poolfstat* [v3.0] package in R (*110*). Analysis tools in *poolfstat* are optimized for pool-seq data and explicitly account for coverage differences among pools through the “effective coverage” of the pool (n_e_) statistic as described by ((*14*, *44*, *111*); eq. 2):

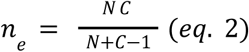

In this context, N is the number of pooled chromosomes and C is the mean read depth of the pool. This formula accounts for the inherent error in allele frequency estimation of Pool-Seq (*112*) within a given pool. We estimated the *F*_ST_ statistic for all six replicates of the F16 lines as a pairwise comparison relative to the VT8_0_1 parental line. We evaluated the signal of the *F*_ST_ statistic at the level of genomic windows based on recombination rates. We report the enrichment analysis of *F*_ST_ values in the top 1% across the genome. We ranked-normalized the *F*_ST_ values and used a binomial enrichment test with the null hypothesis that 1% of SNPs within any given window will have *F*_ST_ values among the top 1% genome-wide. Regions that show higher enrichment of top 1% *F*_ST_ SNPs, relative to our simulations (see above), are candidate haplotypes of adaptive introgression.

### Characterizing footprints of adaptive introgressions using the Fisher’s Exact Test

We identified regions of interest resulting from our adaptive introgression experiment by calculating a Fisher’s Exact Test between the parental and the F16 introgressed lines (both with VT and SK mothers) and the parental Vermont line. We ran this analysis for all alleles in the genome (mAF > 0.05) considering both the allele count and overall coverage to construct the contingency table. To be conservative, we transformed *P*-values into *Q*-values (i.e., adjusted *P*-value) using a false discovery rate transformation (*113*). In the context of this test, we defined a locus of interest as any mutation showing a significant result in a Fisher’s Exact Test within a region identified by our simulations as deviating from the expected pattern of genetic drift.

### Colocalization with other datasets

We assessed whether any of our putative targets of adaptive introgression have been reported in previous studies of ecological adaptation. We did this by comparing the FET *Q*-values reported in our study for the windows of interest (i.e., in 2R and X; see *Results*) with the *P*-values from clinal (*21*) and seasonal (*23*) datasets. For the clinal analysis, we considered that clinal SNPs are those with a clinal *Q-*value less than 0.05. For the seasonal analyses, we used SNPs with seasonal *P*-values less than 0.004. Both thresholds were chosen and used by the authors in their respective papers.

### Linkage disequilibrium and Inversion markers

We calculated levels of linkage disequilibrium among individual mutations using Plink v1.9 (*114*). To assess the levels of linkage between the top candidate in 2R and the cosmopolitan inversion *In(2R)NS*, we estimated levels of linkage between the SNP and the inversion markers identified by (*115*).

### Tests for clinality and seasonality

We used the DEST 2.0 (*34*) dataset (only samples that passed the quality filter recommended by the authors; see table S1 in the reference) in order to study the patterns of genetic variation from the top SNPs emanating from the adaptive introgression and colocalization study. Accordingly, we fit two types of analyses: clinal and seasonal tests. To test for clinality, we calculated the correlation between latitude and the allele frequency at each locus of interest. On the other hand, the seasonal models were performed using a generalized linear model framework using a binomial error structure. For these models, the dependent variable is the estimated allele frequency for each pooled sample, weighted by *n*_e_ (see eq. 2) of the pool. As for the dependent variable, we used the same summary statistics as in the clinal models summaries, yet explicitly summarized across 15, 30, 45, 60, 75, and 90 days prior to the collection date of each sample. We conducted the seasonal test using a framework similar to that of Nunez et al. 2023 (*13*). We report the *P*-*value* of a likelihood ratio test between a two models, one that contains only the year and location of origin (as factors; eq. 3), and another that also includes the environmental variable (eq. 4) as follows:

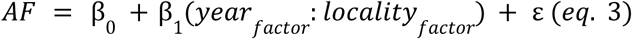

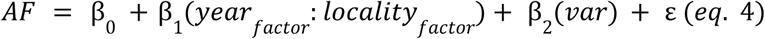

In these equations, β_0_ is the intercept term, β_1_ models the location and year of the collection (as factors), and β_2_ is the environmental variable. We also used 100 random permutations to assess the significance of the models.

### Embryonic heat-shock survival assay in the DGRP

DGRP lines (Table S6) were obtained from the Bloomington Stock Center in Indiana and maintained under common-garden conditions on cornmeal-yeast-molasses medium at 25°C on a 12L:12D light cycle. Following the protocol of Lockwood et al. (2018) (*30*), approx. 50 pairs of 3-5 day-old adult flies were allowed to mate and lay eggs on grape juice agar plates (60 x 15 mm) for 1 h at 25°C. The plates containing embryos were then wrapped in Parafilm and subjected to a 45-minute heat shock in a water bath set to 35°C. As reported in Lockwood et al. (2018) (*30*), this heat treatment produces a heat ramp of approx. +0.6°C min^-1^, which is a rate of temperature increase within the range of heating rates of necrotic fruit in nature (*116*). Following heat exposure, embryos were carefully arranged into 5×4 grids across the surface of the agar with a fine paintbrush. The plates were then returned to 25°C for a 48-hour recovery period. Embryonic survival was quantified as the proportion of hatched embryos observed after the recovery period.

### Gene expression

We conducted RNA-seq to assess the extent to which candidate genes exhibited differential expression based on genetic background and exposure to heat stress. Following the introgression mapping, we subjected each replicate introgression to another round of heat selection (80% mortality, as previously) and isolated three mated females per replicate cross, which were used to establish three isofemale lines. Each line was subsequently inbred with full-sib mating for two generations to reduce the likelihood of lab evolution (*96*).

To assess the gene expression response of embryos to heat stress, and following the protocol of (*75*), we subjected 0-1 h old embryos to an acute temperature treatment of 45 minutes at 25°C or 34°C. For each introgression mapping replicate, we pooled embryos across the three isofemale lines to incorporate the potential effects of genetic variation among isofemale lines. For each parental line (i.e., SK or VT), embryos were collected from the original parental line and thus do not represent pools across separate lines. A total of 100 embryos were pooled for subsequent RNA extraction and sequencing.

Embryos were processed for RNA sequencing following the protocol of (*75*). Embryos were washed with 0.7% NaCl, 0.05% Triton X-100 for 30 seconds, dechorionated for 1 minute with 50% bleach, washed for 30 seconds in RO water, immediately frozen on liquid nitrogen, and stored at -80°C. Samples were thawed on ice and homogenized in 150µL Trizol. We then added an additional 250µL of Trizol, 150µL of chloroform, and 150µL of DEPC-treated water to each sample and centrifuged at 4°C at 12,000g for ten minutes. RNA was precipitated with 375µL 100% isopropanol for 10 minutes at room temperature, then centrifuged for ten minutes at 4°C at 12,000g, and washed with 750µL of 75% EtOH, then centrifuged at 4°C at 7,500g for five minutes. Samples were air dried and then resuspended in 80µL of DEPC-treated water and incubated at 4°C for ten minutes. We re-precipitated samples with 8µL 3M NaOAc and 200µL 100% EtOH and incubated them at -80°C for 30 minutes before centrifuging at 4°C at maximum speed for 30 minutes. After again air drying, samples were resuspended in 18.5µL DEPC-treated water. Total RNA quantity and purity were assessed using a Nanodrop 2000 spectrophotometer (Thermo Scientific). Total RNA integrity was assessed using a 2% agarose electrophoresis gel or a Bioanalyzer 2100 (Agilent Technologies). mRNA was enriched using oligo(dT) beads, then cDNA was synthesized using mRNA templates and random hexamer primers. Sample library quality was assessed by Qubit 2.0 Fluorometer (Thermo Scientific), BioAnalzyzer 2100 (Agilent Technologies), and real time PCR. The libraries were sequenced as 150nt paired-end reads using Novaseq 6000 (S4 Flowcell, Illumina). We checked the quality of paired-end raw sequence reads using FastQC (v. 0.11.7) (*117*). We then trimmed the forward and reverse reads using Trimmomatic (v. 0.38) (*118*) to remove adapter sequences and low quality leading and trailing bases. Reads were aligned to the reference genome (DM6 with ensembl gene annotation v.10) and quantified using salmon (v. 0.14.1) (*119*). We used R (v. 4.2.3) and the package “DESeq2” (v. 1.38.3) to normalize read counts and quantify gene expression (*120*). We note that these transcriptomic data were utilized herein to examine the expression of few candidate genes of interest but are not presented in their entirety. The full scale analysis of these data is beyond the scope of the present study and will be presented in a forthcoming manuscript.

## Acknowledgements

We thank Melissa Pespeni, Reid Brennan, Sara Helms Cahan, Kristi Montooth, Colin Meiklejohn, Thomas O’Leary, Mark C. Bitter, and John Pool for helpful discussions. JCBN acknowledges the Henderson-Harris fellowship program at the University of Vermont for institutional support, and the Vermont Advanced Computing Center (VACC; https://www.uvm.edu/vacc) for providing computational resources that contributed to this publication.

## Data availability

Code for all analyses presented in this paper is available in Github at https://github.com/bllockwo/embryonic_introgression_mapping_VTtropical_2020. The raw DNA reads for the parental and introgression lines are available in the National Center for Biotechnology Information (NCBI), short read archive (SRA), SRR ids: SRR34395643-SRR34395650 (in PRJNA1285949). The raw RNA reads can be found as SRR ids: SRR34364632-SRR34364655 (in PRJNA1285997). Information regarding the DEST data set can be found at (https://dest.bio/), and the pipeline is available in Github at https://github.com/DEST-bio/DESTv2.

