## Supplementary material for "Early life-stage thermal resilience is determined by climate-linked regulatory variation": Fig. S

### Supplementary materials for Early life-stage thermal resilience is determined by climate-linked regulatory variation

✤ Equal contribution
♮Current address: Chulalongkorn University, Bangkok, Thailand
Φ Current address: Dept. of Organismic and Evolutionary Biology, Harvard University, Cambridge, MA, USA

---

**Figures (in this PDF)**
Figure S1: Embryonic heat tolerance ($LT_{80}$) of F16 samples.
Figure S2: Coverage and nucleotide diversity for pooled samples.
Figure S3: Genomic similarity between the F16 samples and the VT and SK parents.
Figure S4: Sliding window analysis for $F_{ST}$ outliers in the introgression test.
Figure S5: Patterns of allele frequency change from Pool-Seq and individual sequencing data.
Figure S6: The top hits in the X chromosome and embryonic phenotypes.
Figure S7: Patterns of allele frequency introgression in the region in the X chromosome.

**Tables (downloadable from the publisher's website)**
Table S1: Sample metadata.
Table S2: Top SNPs in Fisher's exact test.
Table S3: Gene ontology (GO) analyses in 2R.
Table S4: Gene ontology (GO) analyses in X.
Table S5: Gene ontology (GO) analyses genome-wide.
Table S6: Embryonic heat shock survival data in the DGRP
Table S7: Developmental gene expression data for genes related to the top SNP in 2R.
Table S8: Regulatory SNP annotations associated with *SP70* and *sog*.

**Texts (in this PDF)**
Extended Materials and Methods
Text S1: Details on the GO enrichment analysis.
Text S2: The evolutionary context of our top SNPs in 2R and X.

**Figure S1:** Embryonic heat tolerance ($LT_{80}$) of F16 samples.

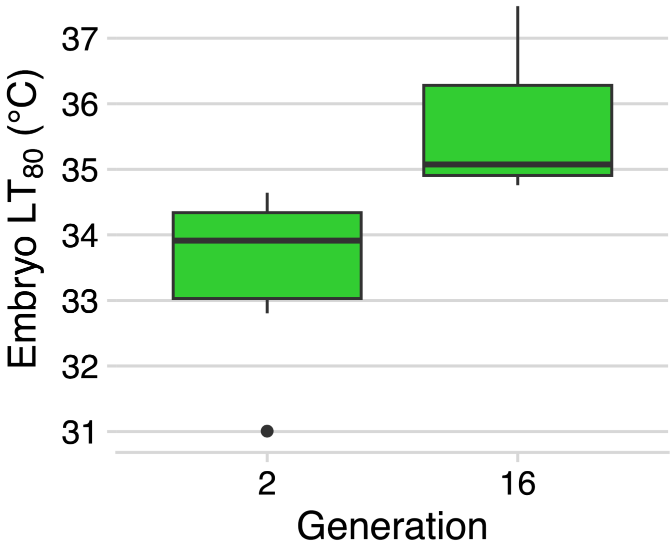


Embryonic heat tolerance ($LT_{80}$) estimated from the exposure (1 hour at the indicated temperature) that induced 80% mortality among six replicate F2 and F16 introgression lines.

**Figure S2:** Coverage and nucleotide diversity for pooled samples.

**A**

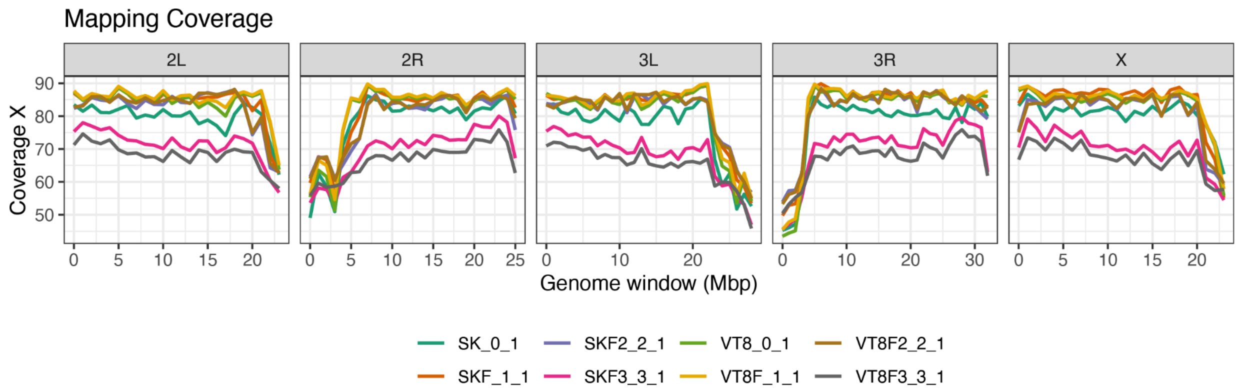


**B**

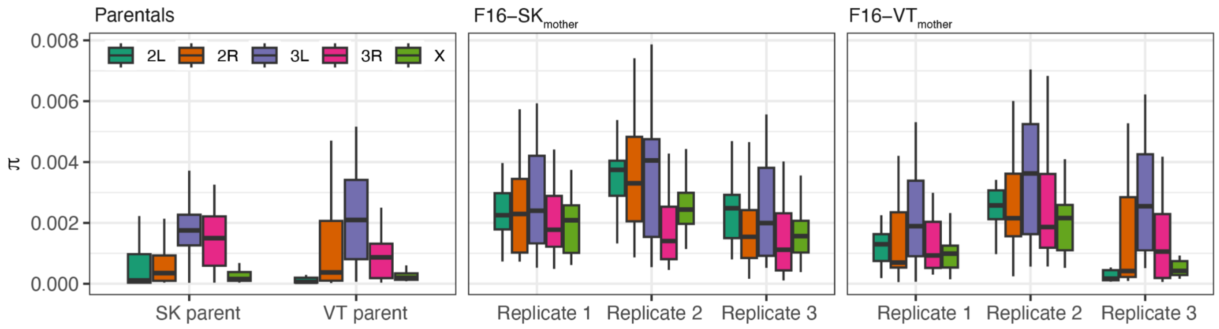


**(A)** Coverage across chromosomes of *D. melanogaster* for all pools. To investigate the genomic basis of this response, we performed pooled whole-genome sequencing on six replicate F16 introgressed populations: in three, VT was the maternal parent, and in the other three, SK was the maternal parent. Sequencing was also performed on the original VT and SK parental strains. Sequencing results of both the parental pools and the six F16 introgression pools have deep levels of coverage (mean = 87.6X, sd = 14.1X) and high mapping quality scores (mean = 56.6, sd = 0.249). Considered individually, the parental lines and the replicate pools 1 and 2 for F16 crosses have similar coverages (78X-82X). Pools from the third F16 replicate have slightly lower, yet deep, levels of coverage (66X-69X). Given pools were sequenced on individual flow cells, these coverage differences suggest a mild technical effect for the third replicate of our crosses. Yet, this is not a concern for downstream analyses since our analytical tools account for coverage differences by design (see materials and methods). The estimated levels of PCR and optical duplicates filtered during mapping was ~0.25% (sd = 0.03%). Combined, these results indicate that our sequencing efforts yielded high-quality data. The panels show the mean coverage across each chromosome. The colors, indicated in the legend, are each replicate pool. Parental lines are labeled as “0_1” (e.g., SK_0_1 and VT8_0_1), and replicates of the F16s are labeled as “n_1” where n is the replicates (e.g. , 1, 2, 3). **(B)** Levels of nucleotide diversity ($\pi$) across chromosomes for parental lines and all F16 pools, contingent on whether SK or VT was used as the mother in the cross. After applying a 5% minimum threshold for missing data, we discovered 865,020 SNPs segregating among all pools. Of these, 159,641 are found in 2L, 205,515 in 2R, 187,745 in 3L, 203,632 in 3R, and 108,859 in X. Analyses of nucleotide diversity (π) showed lower π in our parental lines (ANOVA; $F_{2,1085}$ = 63.12, $P = 2.0\text{x}10^{-16}$) relative to our introgressed lines. For example, parental inbred lines in VT and SK had $\pi$ levels ranging from 0.10-0.11. In contrast, F16 lines had doubled the level of π (i.e., 0.239 and 0.186; for the F16 initiated with SK-mothers and VT-mothers, respectively). These data revealed two salient results: first, the SK and VT parental lines harbor residual heterozygosity. Second, 7 generations of introgression did not return π to the original levels of the parental VT lines, a likely byproduct of the segregating residual heterozygosity from the initial SK x VT cross.

**Figure S3:** Genomic similarity between the F16 samples and the VT and SK parents.

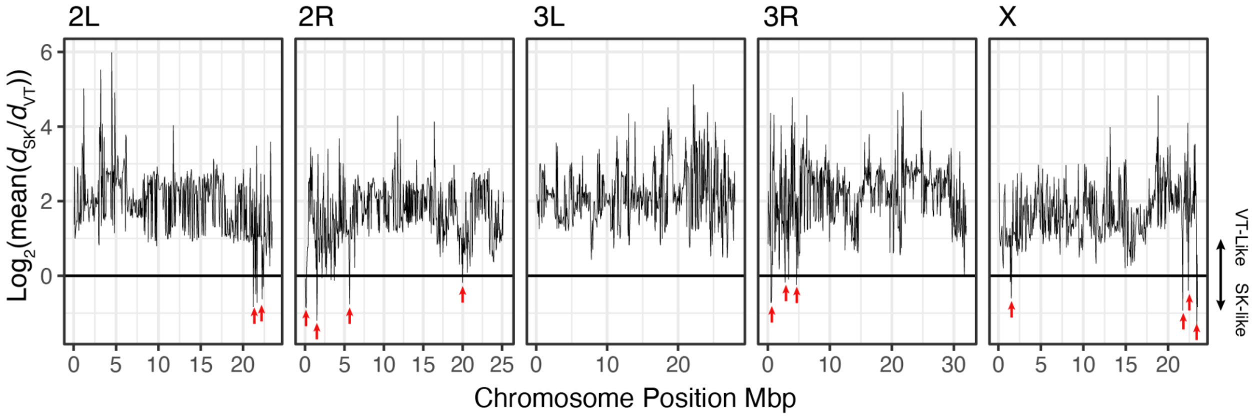


To investigate whether particular regions of the F16 genomes show differential patterns of introgression, relative to the parental lines, we estimated local PCAs using a sliding window approach (window size = 0.1 Mb; step = 50 Kb). We then estimated the mean euclidean distance (i.e., $d$), using PCs 1 and 2, across F16 pools relative to each parent (i.e., $d_{SK}$ and $d_{VT}$). Accordingly, the ratio of $d_{SK}$ and $d_{VT}$ is a proxy for genetic distance, whereby $Log_2(d_{SK}/d_{VT}) > 0$ indicates higher similarity to the VT parent and $Log_2(d_{SK}/d_{VT}) < 0$ indicates higher similarity to the SK parent.

**Figure S4:** Sliding window analysis for $F_{ST}$ outliers in the introgression test.

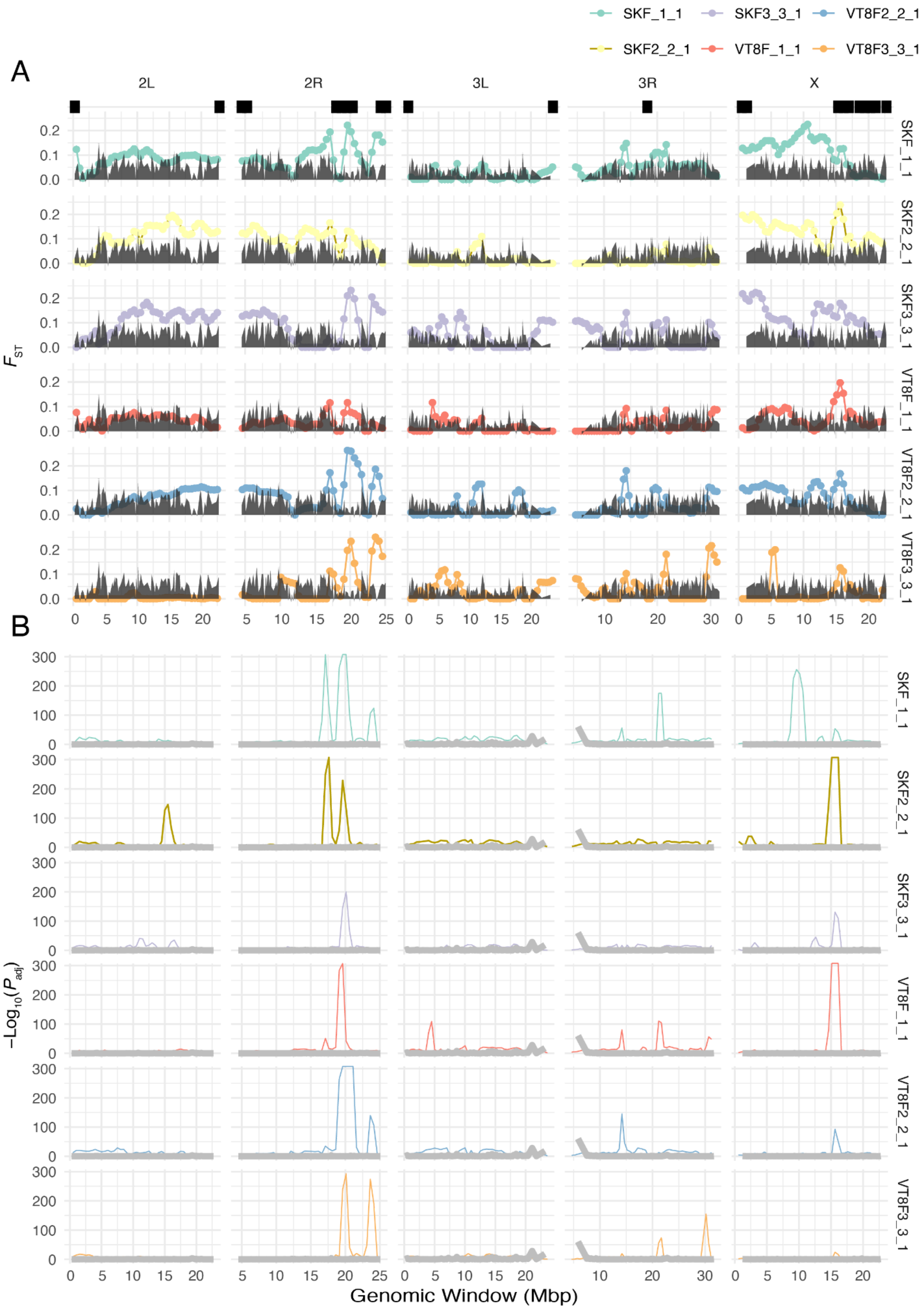


**(A)** Sliding window analysis for $F_{ST}$ outliers across our introgression crosses relative to neutral introgression simulations across all chromosomes. **(B)** Sliding window analysis for the enrichment of $F_{ST}$ in the top 1% of the distribution across the genome.

**Figure S5:** Patterns of allele frequency change from Pool-Seq and individual sequencing data.

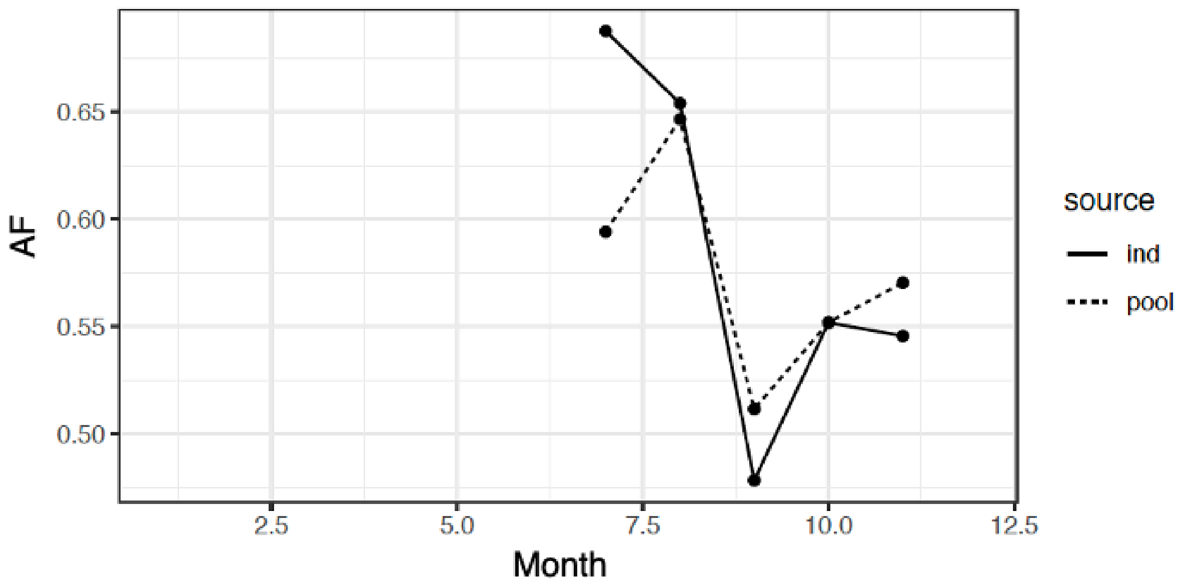


Patterns of allele frequency change from Pool-Seq (dashed lines) and individual (solid lines) sequencing data. Allele frequency estimates from pooled sequencing (DEST Virginia samples) are shown alongside individually sequenced flies from Nunez et al. (2024), collected at the same site in 2016. Patterns of allele frequency change are concordant between datasets (Granger Causality Test, $P = 0.032$).

**Figure S6:** The top hits in the X chromosome and embryonic phenotypes.

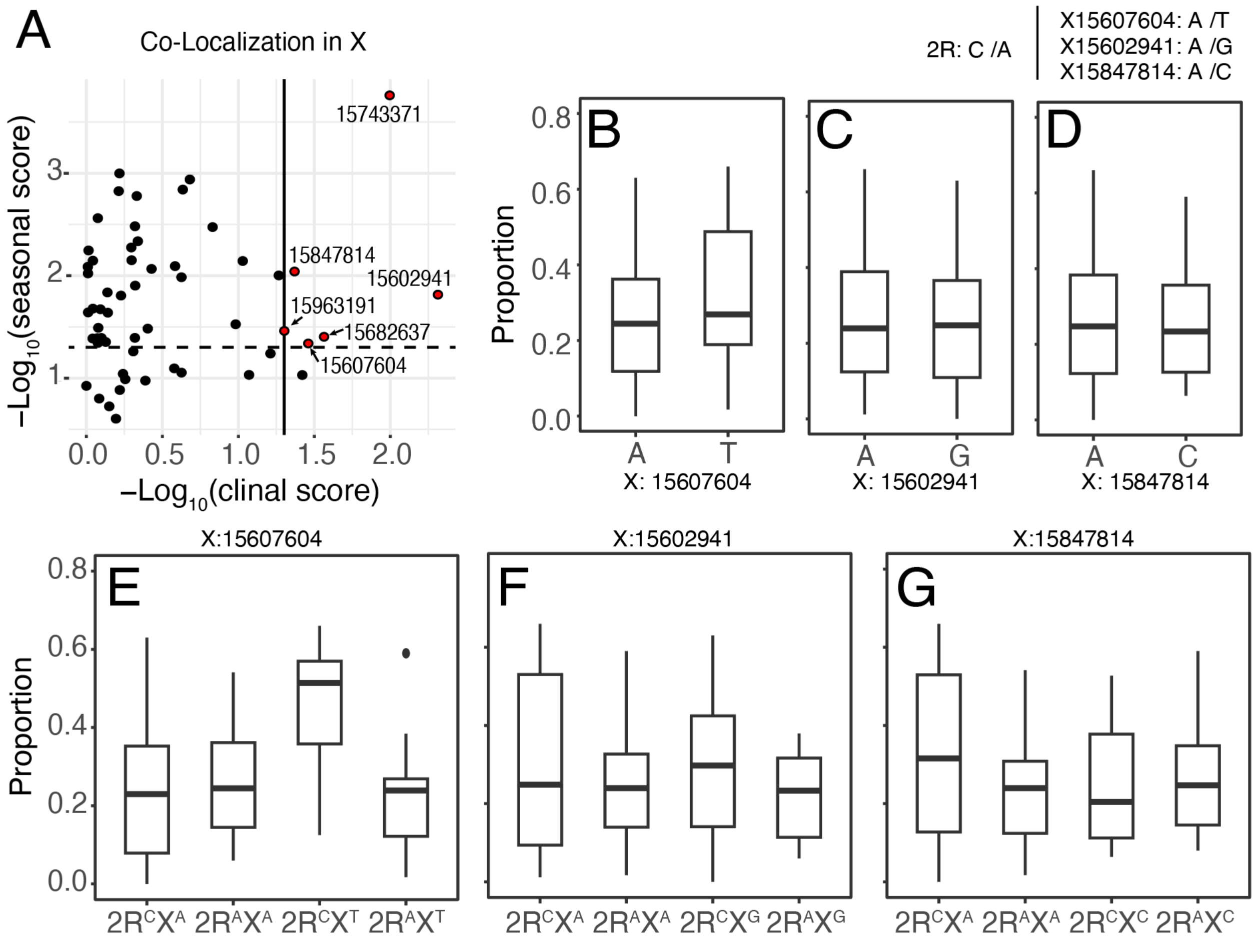


**(A)** SNPs that are jointly FET outliers, seasonal outliers, and clinal outliers in the window of interest of the X chromosome. **(B-D)** Embryonic survival for genotypes across the top three SNPs in the X chromosomes. **(E-G)** Embryonic survival for genotype combination across the top hit in 2R and the three top SNPs in the X chromosomes. The only combination with a significant effect, $2R^{C}X^{T}$, shown in panel E, is the same combination highlighted in the main text as the "tropical genotype."

**Figure S7:** Patterns of allele frequency introgression in the region in the X chromosome.

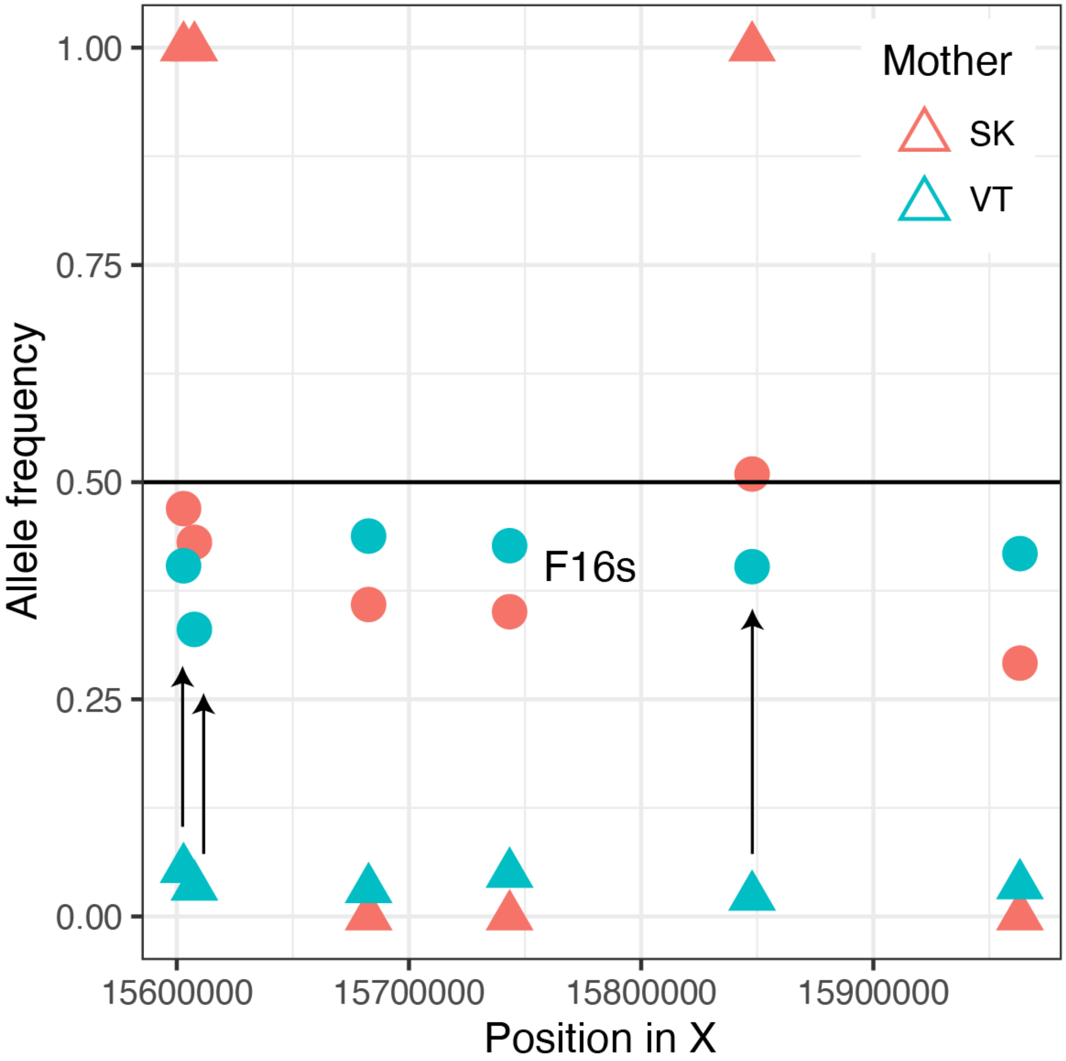


Patterns of allele frequency introgression in the region of interest of the X-chromosome. The red triangles indicate the allele frequency in the SK background. The blue triangles indicate the allele frequency in the VT background. The circles indicate the allele frequency of the F16 offspring, the color of the circle indicates the background of the mother. SNPs showing patterns consistent with adaptive introgression (i.e., towards the SK background) are indicated with arrows.
