## Supplementary material for "Early life-stage thermal resilience is determined by climate-linked regulatory variation": Text S1

***Text S1: Gene Ontology Enrichment analyses***

We calculated GO enrichment scores using the outlier SNPs, derived from the synthetic pool FET, harbored inside of the windows of interest in 2R and X. For the case of the window in 2R, we observe one GO term with a *Q*-value threshold below 0.1. The term (GO:0005549), is associated with odorant binding functions (*Q* = 0.081) and it is driven by the presence of outlier SNPs at three paralogs of the Odorant-binding proteins 56 and 57 (*Obp56/57*) gene (*Obp56d*, *Obp57g*, *Obp57d*). Two other GO terms in the enrichment are related to heat shock protein (*Hsp70* and *Hsp90*) binding, GO:0030544 and GO:0051879. These terms have low *P*-values (*P*-value = 0.003, *P*-value = 0.005, respectively) but do not pass *Q*-value filtering (*Q*-value > 0.1) in our analysis. Yet, these terms appear in the analysis due to the presence of the *CG15120* gene (FBgn0034454). A gene predicted to play a role in *Hsp70*, *Hsp90*, and ubiquitin protein ligase binding activity.

For the window in the X chromosome, we observed 10 GO terms enriched with a *Q*-value threshold below 0.1. Two of these terms are associated with the regulation of euchromatin (GO:0005719, GO:0000791; *Q*-values of 0.013 and 0.062, respectively) and are driven by two genes: Myb oncogene-like (*Myb*) and DNA oxidative demethylase (*AlkB*). Six are related to t-RNA ligase activity, tRNA aminoacylation, amino acid activation, as well as other functions related to carbon-oxygen bonds (GO:0016876, GO:0016875, GO:0006418, GO:0004812, GO:0043039, GO:0043038; *Q*-values = 0.062 throughout). Two genes primarily drive the enrichment of these terms, *CG8097*, which encodes the arginine--tRNA ligase enzyme, and the Arginyl-tRNA synthetase (*ArgRS*) gene. The last two terms are related to centriole assembly and replication (GO:0007099, GO:0098534; *Q*-values = 0.068 for both). These terms are driven by SNPs found in the aforementioned *Myb*, as well as of the Chaperonin containing TCP1 subunit 6 (*CCT6*) gene.
