## Supplementary material for "Early life-stage thermal resilience is determined by climate-linked regulatory variation": Text S2

***Text S2: The evolutionary context of our top SNPs in 2R and X***

To better understand the findings from our adaptive introgression experiment, we integrated findings from ecological genomics analyses using the DEST dataset. Through this approach, we identified two high-confidence SNPs within our focal regions (2R:20,551,633 and X:15,607,604). In the case of 2R, the tropical allele ("C") is the ancestral allele relative to four outgroup species (*D. simulans, D. yakuba, D. sechellia, and D. erecta*) and it is the major allele in Africa. For the case of X, the tropical ("T") is the ancestral allele relative to *D. simulans,* and *D. sechellia*; notably, *D. yakuba* shows a different allele altogether ("G").

While both mutations showed a response to selection in our experiment, we did not detect any meaningful linkage disequilibrium (LD) between them in VA ($r^2_{\text{X-2R SNPs}}$ = 0.0008) or in the DGRP panel ($r^2_{\text{X-2R SNPs}}$ = $1.232 \times 10^{-5}$). For the particular case of our 2R candidate, given the close physical proximity of our top SNP to the cosmopolitan inversion *In(2R)NS*, we investigated the extent of potential linkage between these loci. To this end, we calculated LD within a genomic window of approximately $10^7$ bp in the DGRP panel, including a set of established inversion markers for *In(2R)NS* (see materials and methods). The results showed that, despite its physical proximity to the inversion, the top SNP in 2R segregates independently of *In(2R)NS* (**Fig. TS2-1**).

We also inferred the relative ages of the mutation and the inversion using a phased dataset from wild Virginia populations. Our results suggest that both the 2R and X mutations are old ($\text{TMRCA}_{\text{2R SNP}}$= 214,384.8 ya; $\text{TMRCA}_{\text{X SNP}}$= 151,074.8 ya), predating the most recent out-of-Africa migration by at least 140,000-200,000 years. In the particular case of 2R, the top SNP emerged roughly three times earlier than the *In(2R)NS* inversion (Mean $\text{TMRCA}_{\text{inv. markers}}$ = 62,303.9 ya).

**Figure TS2-1:**

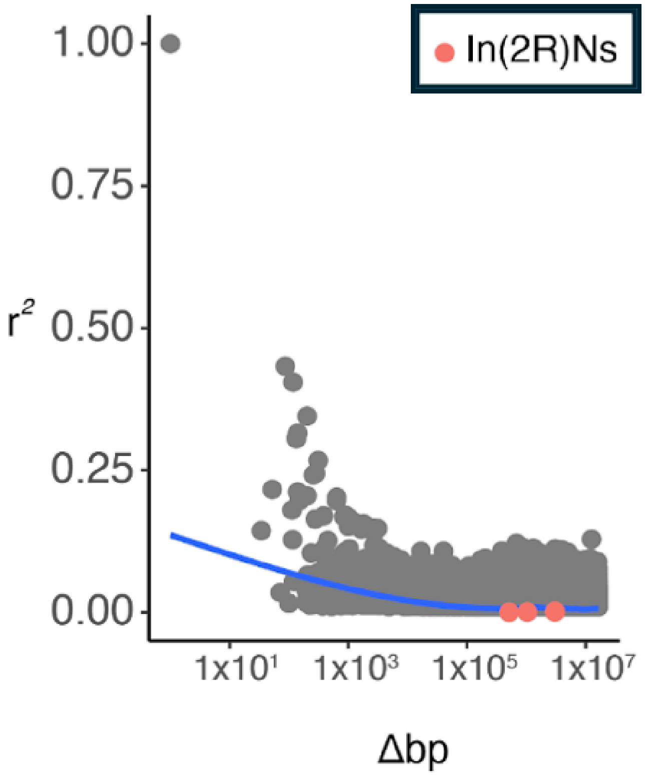


Patterns of linkage disequilibrium ($r^2$) in the window of interest in chromosome 2R, in relationship to the top SNP in 2R. The in(2R)NS markers are shown in red.
